# The crystal structure of TRPM2 MHR1/2 domain reveals a conserved Zn^2+^-binding domain essential for ligand binding and activity

**DOI:** 10.1101/2022.01.04.474898

**Authors:** Simon Sander, Ellen Gattkowski, Jelena Pick, Ralf Fliegert, Henning Tidow

**Author notes:** Corresponding author: Henning Tidow, University of Hamburg, Department of Chemistry, Institute for Biochemistry and Molecular Biology, Luruper Chaussee 149, D-22761 Hamburg, Germany.

## Abstract

Transient receptor potential melastatin 2 (TRPM2) is a Ca^2+^-permeable, non-selective cation channel involved in diverse physiological processes such as immune response, apoptosis and body temperature sensing. TRPM2 is activated by ADP-ribose (ADPR) and 2′-deoxy-ADPR in a Ca^2+^-dependent manner. While two species-specific binding sites exist for ADPR, a binding site for 2′-deoxy-ADPR is not known yet. Here, we report the crystal structure of the MHR1/2 domain of TRPM2 from zebrafish (*Danio rerio*) and show binding of both ligands to this domain. We identified a so-far unrecognized Zn^2+^-binding domain that was not resolved in previous cryo-EM structures and that is conserved in most TRPM channels. In combination with patch clamp experiments, we comprehensively characterize the effect of the Zn^2+^-binding domain on TRPM2 activation. Our results provide insight into a conserved structural element essential for channel activity.

## Introduction

Members of the melastatin subfamily of transient receptor potential channels (TRPM channels) are widely expressed and contribute to cellular Ca^2+^ signaling either directly or indirectly. They play an important role in physiological processes such as temperature sensing and regulation (TRPM8 (McKemy et al., 2002), TRPM2 (Siemens et al., 2016; Tan & McNaughton, 2016)), the immune response (Schmitz & Perraud, 2005), Mg^2+^ homeostasis (TRPM6 / TRPM7 (Montell, 2003), taste sensing (TRPM5, (Pérez et al., 2002)), the response to oxidative stress (Simon et al., 2013) and apoptosis (McNulty & Fonfria, 2005). Many diseases are linked to TRPM proteins (Jimenez et al., 2020) explaining the increased attention they receive as potential drug targets (Abriel et al., 2012; Sun et al., 2015; Vennekens et al., 2018). The activity of TRPM channels is regulated by various influences such as voltage and temperature as well as changes in concentrations of small molecules, ions or lipids (reviewed in (Huang et al., 2020)). Structurally, all TRPM family members share a common core architecture: N-terminal TRPM homology regions (MHR1-4), six transmembrane helices, a TRP helix and a C-terminal coiled-coil domain (Huang et al., 2020).

The Ca^2+^-permeable, non-selective cation channel TRPM2, which is involved in cell death (Hecquet et al., 2014), diverse immune cell functions (Knowles et al., 2011) and the control of body temperature (Siemens et al., 2016; Tan & McNaughton, 2016) harbors a unique C-terminal domain. The name of this domain (NUDT9-H) arises from the homology to the Nudix box enzyme NUDT9, a soluble pyrophosphatase, that hydrolyzes adenosine 5’-diphosphoribose (ADPR). Due to this homology the NUDT9-H domain was predicted to contain a binding pocket for ADPR, the endogenous ligand of TRPM2 (Perraud et al., 2001). Although human TRPM2 does not possess ADPR hydrolase activity (Iordanov et al., 2016), the recently determined structure of human TRPM2 illustrated that the NUDT9-H domain in fact binds ADPR (Huang et al., 2019). However, the cryo-EM structure of TRPM2 from zebra fish revealed a distinct ADPR-binding site in the N-terminal MHR1/2 domain (Huang et al., 2018). Since this site is conserved across TRPM2 from different species including humans, human TRPM2 harbours two distinct ADPR-binding pockets (Huang et al., 2018, 2019; Wang et al., 2018). The current view of gating of TRPM2 by ADPR is, that upon ADPR binding in the N-terminal domain, the clamshell-like shape of the MHR1/2 domain closes, inducing a rotation. This rotation and binding of Ca^2+^ at the membrane-cytosol interface as well as binding of a second ADPR molecule within the NUDT9-H domain leads to further conformational changes in the tetrameric channel provoking an activated state (Huang et al., 2018, 2019; Yin et al., 2019).

The cellular nucleotide ADPR is a metabolite of NAD (nicotinamide adenine dinucleotide) and can arise from the hydrolysis of NAD by the multifunctional enzyme CD38 (Howard et al., 1993; Zocchi et al., 1993). Other sources are proteins that were marked with polymeric ADPR residues by the poly-ADPR polymerase (PARP). The hydrolysis of poly-ADPR and the formation of free monomeric ADPR is catalyzed by the poly-ADPR glycohydrolase (PARG) and the terminal ADPR protein glycohydrolase (TARG) (Nikiforov et al., 2015). The PARP/PARG pathway is triggered by oxidative stress explaining the connection between the oxidative stress response and TRPM2 (Buelow et al., 2008). Another TRPM2 activator is 2′-deoxy-ADPR, which proved to be a more effective TRPM2 agonist than ADPR. 2′-deoxy-APDR could as well be detected endogenously and thus may act as second messenger (Fliegert et al., 2017).

Like a number of other Ca^2+^ channels, the human TRPM channels TRPM3, TRPM6 and TRPM7 as well as the homologous single TRPM channels from Drosophila melanogaster have been shown to be permeable to Zn^2+^ and to mediate Zn^2+^ entry upon activation (Bouron & Oberwinkler, 2014). While TRPM2 is inhibited by high extracellular Zn^2+^ (Yang et al., 2011), it has been shown that the cytosolic Zn^2+^ concentration in TRPM2 expressing cells increases upon activation of a photoactivatable ADPR analogue (Yu et al., 2012). This indicates that TRPM channels contribute to the cellular Zn^2+^ homeostasis. Zn^2+^ is an essential trace metal and required as cofactor for some enzymes and structural components of a large number of proteins (reviewed in (Kambe et al., 2015)). It has also been considered to exert signaling function and act as second messenger (Fukada et al., 2011). Similar to Ca^2+^, high concentrations of Zn^2+^ are cytotoxic and contribute to excitotoxicity (Granzotto et al., 2020) and other pathological processes. Investigating the role of TRPM2 in cell death of microglial cells a recent study showed that extracellular Zn^2+^ could also activate TRPM2 through multiple steps, involving reactive oxygen species (ROS) that induce PARP/PARG (Mortadza et al., 2017). While the mechanisms remain unknown, there is evidence that also a change in the intracellular Zn^2+^ concentration influences the oxidative stress response and TRPM2 (Abuarab et al., 2017; Ye et al., 2014).

Another member of the TRP family, TRPC5, comprises an intracellular Zn^2+^-binding motif that is conserved within TRPC channels and suggests a direct modulatory role for Zn^2+^ ions on these related proteins (Park et al., 2019; Wright et al., 2020).

In order to improve our understanding of the regulatory role of ADPR and 2′-deoxy-ADPR on the TRPM2 channel, we biophysically characterized ligand binding to the N-terminal MHR1/2 domain of zebrafish TRPM2. The crystal structure of this domain revealed a novel Zn^2+^-binding motif that is conserved within the TRPM family and is essential for channel activity.

## Results and Discussion

### Biophysical characterization of ADPR and 2’-deoxy-ADPR binding to drMHR1/2

The N-terminal MHR1/2 domain of TRPM2 contains a binding site for the channel activating ligand ADPR, which has been identified in TRPM2 cryo-EM structures from human and zebrafish (Huang et al., 2018, 2019). Whether the TRPM2 superagonist 2′-deoxy-ADPR (Fliegert et al., 2017) binds to the same site is currently unknown. In order to investigate the thermodynamic parameters of ligand binding to MHR1/2 and to confirm the structural as well as functional integrity of our sample, we compared binding of ADPR and 2′-deoxy-ADPR to the isolated drMHR1/2 domain using several biophysical techniques.

Isothermal titration calorimetry (ITC) revealed endothermic binding in the low micromolar range for both ligands with 2′-deoxy-ADPR showing a slightly tighter binding than ADPR (Fig. 1A). To investigate whether ligand binding has an effect on the stability of the protein, we used nano differential scanning fluorimetry (nDSF) titrations. We found that ADPR as well as 2′-deoxy-ADPR both concentration-dependently stabilize the drMHR1/2 domain (Fig. 1B).

**Figure 1.**
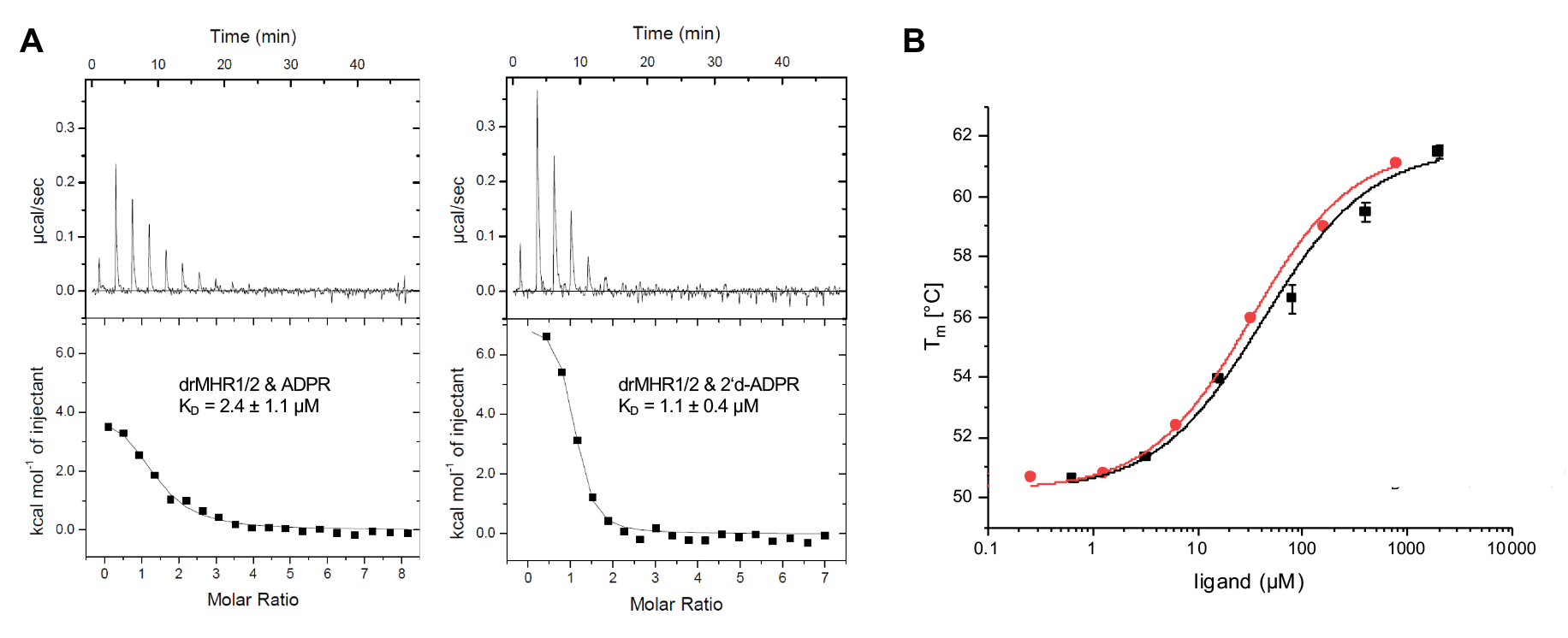
Biophysical characterization of ADPR and 2′-deoxy-ADPR binding to MHR1/2. A) Binding of drMHR1/2 and ADPR / 2’-deoxy-ADPR as measured by isothermal titration calorimetry (ITC). K_D_ values are averages of triplicate experiments. B) Binding of drMHR1/2 and ADPR / 2’-deoxy-ADPR shown by shift of melting temperature observed by differential scanning fluorimetry (nDSF).

### Crystal structure of TRPM2 MHR1/2 domain

The TRPM2 MHR1/2 domain plays a crucial role in the regulation of the full-length channel (Huang et al., 2018, 2019; Tóth et al., 2020). We determined the crystal structure of the zebrafish (*Danio rerio*) TRPM2 MHR1/2 domain (drMHR1/2). Two molecules were present in the asymmetric unit, with crystal contacts generated through the exposed loops. The structure was determined using Se-SAD phasing to 2.0 Å resolution. We could resolve almost an entire MHR1/2 domain with unambiguous side chain information (residues 33-418, with 9 residues missing in the loop comprising residues 201-209). The overall structure (Fig. 2A) shows a bi-lobed clamshell-like shape and superimposes well with the previously published cryo-EM structure of the full-length zebrafish TRPM2 channel (PDB: 6DRK) (Huang et al., 2018) (Figs. 2B-C).

**Figure 2.**
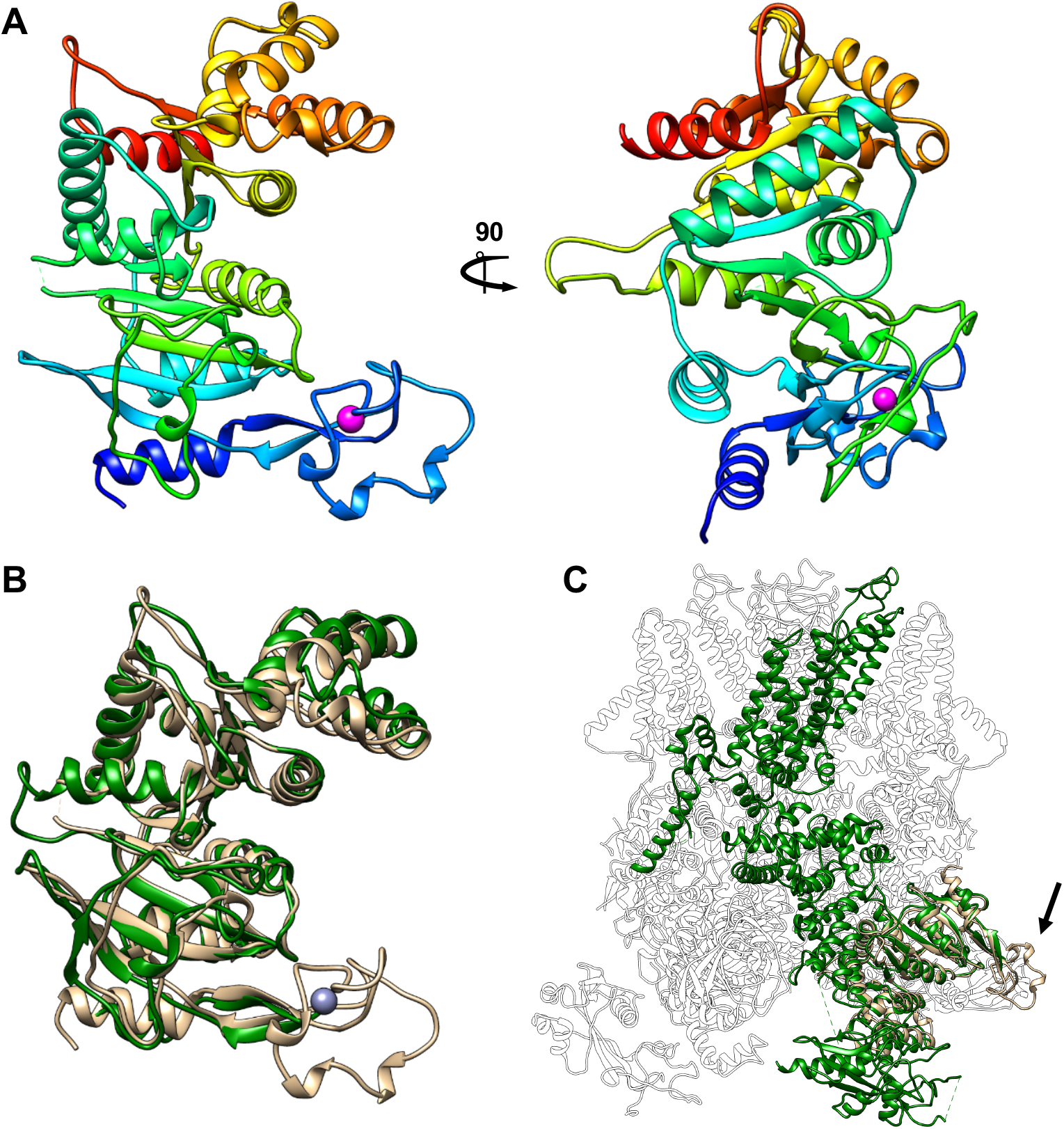
Crystal structure of drMHR1/2. A) The overall fold of drMHR1/2 shows a bi-lobed clamshell-like structure. The structure is colour-coded from blue (N-terminus) to red (C-terminus). Zn^2+^ ion is shown as magenta sphere. B) Overlay of the drMHR1/2 crystal structure (wheat) with the MHR1/2 domain from a drTRPM2 cryo-EM structure (green) (PDB: 6DRK). C) Superposition of the drMHR1/2 crystal structure (wheat) on the tetrameric drTRPM2 cryo-EM structure (white with one monomer in green), indication of the location of the novel Zn^2+^-binding domain (PDB: 6DRK).

While we failed to crystallize drMHR1/2 in complex with ADPR or 2′-deoxy-ADPR, comparison with the horseshoe binding mode of ADPR as observed in the cryo-EM structure of ADPR-bound drTRPM2 (PDB: 6DRJ) indicates that the 2′ hydroxyl group which distinguishes ADPR and 2′-deoxy-ADPR is solvent-exposed in the complex and thus does not contribute to the binding (Huang et al., 2018). This is in agreement with the similar binding parameters obtained for these two ligands by ITC and nDSF.

### Identification and characterization of a conserved Zn^2+^-binding domain

The high resolution of the drMHR1/2 structure allowed unambiguous model building and identified a domain (residues 53-95), located between β1 and β2, which was not resolved in the lower resolution cryo-EM structure of drTRPM2 (Huang et al., 2018). Surprisingly, this novel domain revealed clear electron density for an ion that is coordinated by three cysteines and one histidine (Fig. 3A). The interacting residues C53, C65, C67, and H74 coordinate the ion tetrahedrally (Figure 3B) with geometry and bond length typical for Zn^2+^ ion coordination (according to *CheckMyMetal* server, (Zheng et al., 2017)). An X-ray fluorescence energy scan of the crystal near the zinc absorption K edge (9.6586 keV) unambiguously confirmed the presence of a Zn^2+^ ion (Fig. 3C).

**Figure 3.**
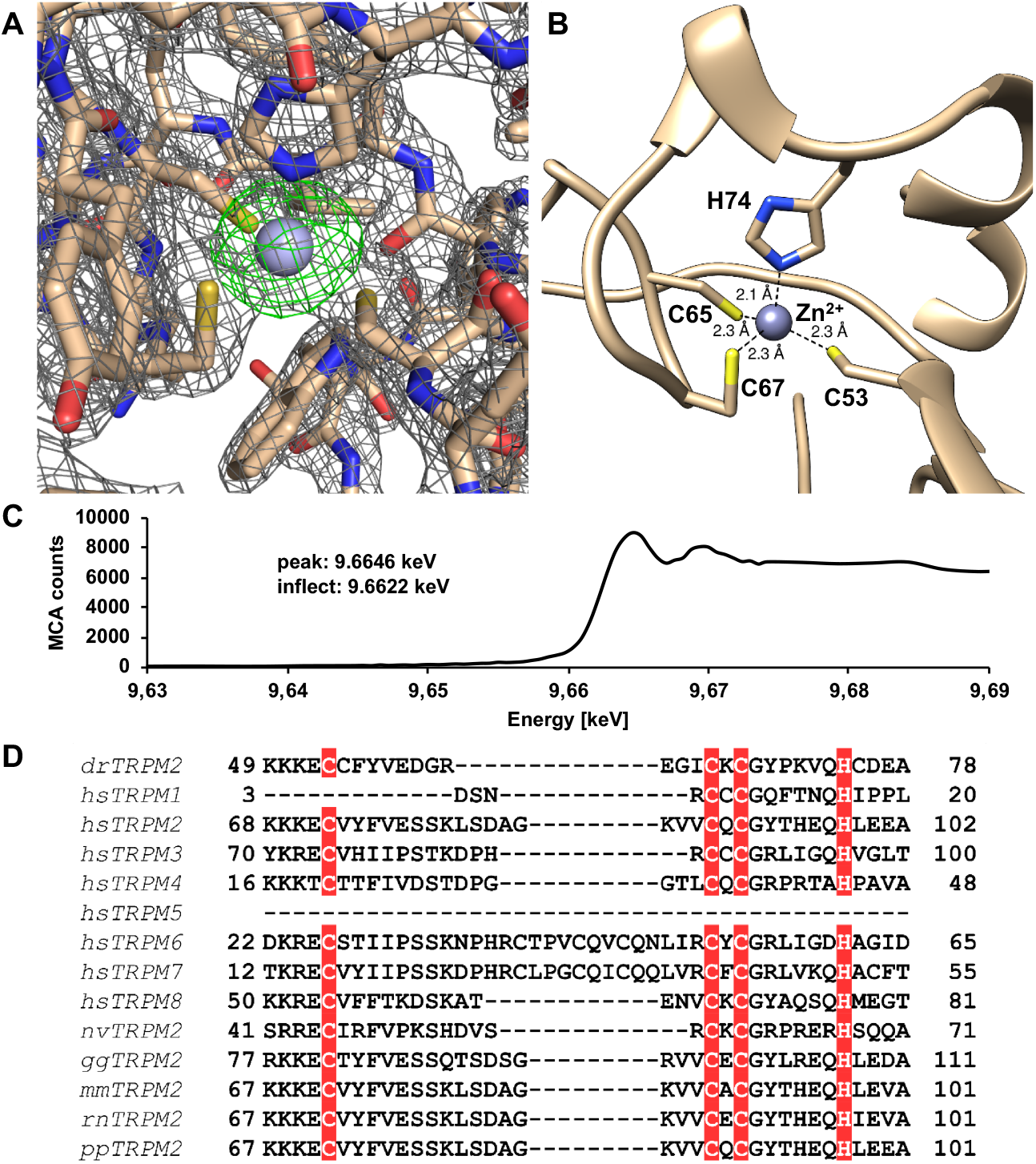
Novel Zn^2+^-binding domain in drMHR1/2 that is conserved in TRPM channels. A) Electron density of the novel Zn^2+^-binding site (grey mesh) and PHENIX POLDER electron density after omission of the Zn^2+^ ion (green mesh) with the atomic model (wheat) and the Zn^2+^ ion (grey). B) Detailed structure view of the Zn^2+^-binding domain with a tetrahedral coordination of the Zn^2+^ ion by C53, C65, C67, H77. C) X-ray energy scan of the fluorescence emitted by the sample near the zinc absorption K edge (9.6586 keV) confirming ion identity.D) Multiple sequence alignment of novel Zn^2+^-binding domain in TRPM channels. Conservation of the four Zn^2+^-coordinating residues (marked) in the N-terminus of most TRPM channels. Alignment of all human (*homo sapiens*) TRPM channels and TRPM2 from sea starlet anemone (*Nematostella vectensis*), chicken (*Gallus gallus*), mouse (*mus musculus*), rat (*Rattus norvegicus*), chimpanzee (*Pan pansicus*), zebrafish (*Danio rerio*).

A multiple sequence alignment revealed that the four residues forming the Zn^2+^-binding domain (C53, C65, C67, H74) are conserved between TRPM2 orthologues from different species and most human TRPM members (Fig. 3D). This strict evolutionary conservation from invertebrates to mammals strongly indicates that the novel Zn^2+^-binding domain is physiologically relevant. As the Zn^2+^-domain with its adjacent ß-stem makes extensive interactions with the remaining MHR1/2 domain we expect a stabilizing effect (Fig. 5). Indeed, we could show its importance for protein integrity/stability by mutating two of the cysteine residues to alanine (C65A and C67A). The mutant drMHR1/2 sequence was recombinantly expressed in *E. coli* but the protein seemed to be insoluble and thus probably incorrectly folded (Suppl. Fig. 1).

### The Zn^2+^-binding domain is required for TRPM2 function / ion conduction

We next set out to investigate the role of the Zn^2+^-binding domain in the context of full-length TRPM2. Full-length human TRPM2 containing equivalent mutations of the Zn^2+^-coordinating residues (in human: C89A and C91A) could be successfully expressed in transfected HEK293 cells and localized to the cell surface, albeit with reduced expression levels compared to the wild-type protein (Fig. 4A). Whole-cell patch clamp measurements revealed that the Zn^2+^-binding site is required for channel activity as the TRPM2 mutants on the cell surface with mutations of Zn^2+^-coordinating residues did not invoke a current upon infusion of ADPR like the wild type protein (Fig. 4B).

**Figure 4.**
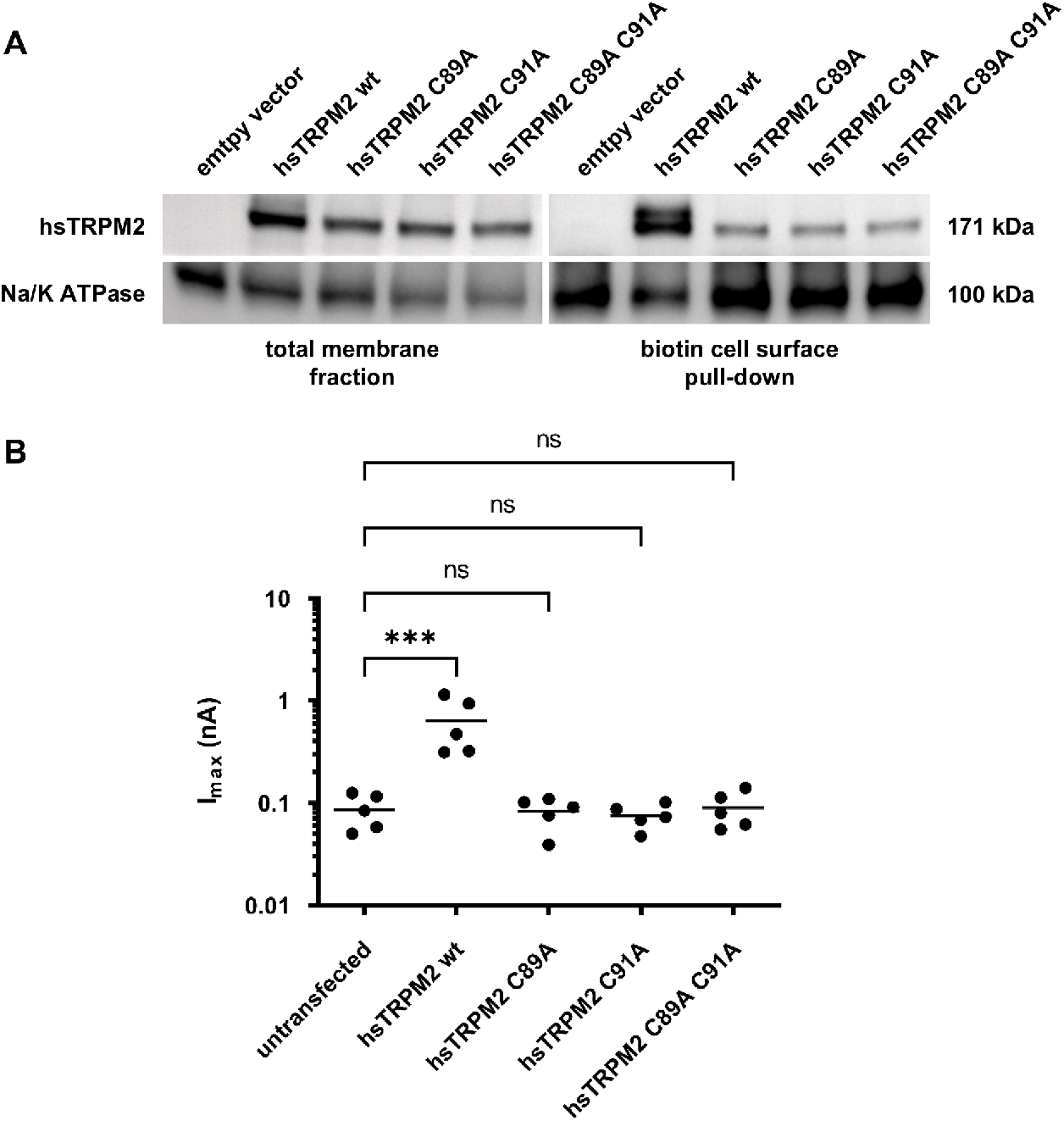
The novel Zn^2+^-binding site in the MHR1/2 domain is important for protein integrity and TRPM2 channel activity. A) Cell surface biotinylation assay with Zn^2+^-binding site hsTRPM2 mutants proving the importance of the motif. The expression level of the mutants is lower on the cell surface as well as in the total membrane fraction. B) Whole-cell patch clamp measurements showing that the Zn^2+^-binding site is also important for channel activity. The remaining fraction of mutant protein on the cell surface of HEK293 wt cells transfected with hsTRPM2 variants does not invoke a current upon infusion of 100 µM ADPR via the patch pipette.

Taken together, we could show that the novel motif is not only crucial for correct protein folding but also plays a role in TRPM2 channel activity. Since the channel with mutations within the motif could not be activated by ADPR, an endogenous ligand of TRPM2, we postulate that the presence of an intact Zn^2+^-domain leads to correct positioning of loop 263-273 (287-297 in hsTRPM2). This loop contains the conserved tyrosine residue Y271 (Y295 in human) which stacks with the adenine moiety of ADPR (see PDB: 6DRJ, (Huang et al., 2018)). In this model the stabilization and loop positioning caused by the Zn^2+^-domain primes the MHR1/2 domain for ligand binding (Fig. 5).

**Figure 5.**
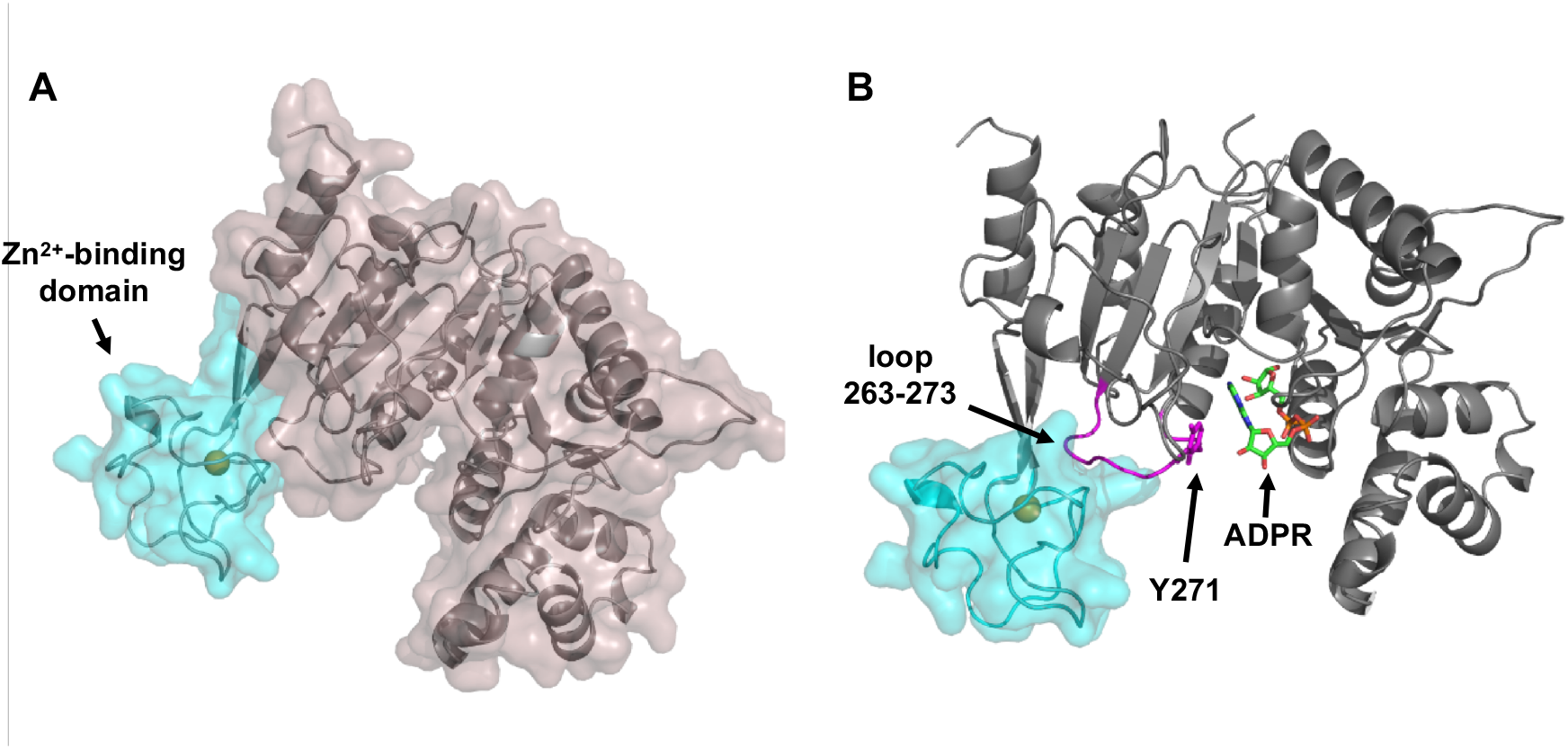
Importance of Zn-binding domain for MHR1/2 stability and ADPR binding. A) Zn^2+^-binding domain (cyan) makes substantial interactions with the remaining part of the MHR1/2 domain (grey). B) Presence of Zn^2+^-binding domain in drMHR1/2 leads to correct positioning of loop 263-273 (magenta) containing the conserved tyrosine residue Y271, which stacks with the adenine moiety of ADPR (see PDB:6DRJ, (Huang et al., 2018)). According to this model the stabilization and loop positioning caused by the Zn^2+^-domain primes the MHR1/2 domain for ligand binding.

Since the Zn^2+^-binding site is structurally exposed to the cytosol it could be possible that it not only provides structural integrity but is furthermore involved in signal transduction with Zn^2+^ as a second messenger. The Zn^2+^ ion could be released from the Cys_3_-His_1_ coordination under oxidative stress or conditions of redox signaling through aldehydes, which has been shown before for other Zn^2+^-binding proteins (Hao & Maret, 2006). Redox signaling pathways that involve possible disulfide bridges connecting C65 and C67 as well as C59 and C80 would lead to reversible binding processes of the Zn^2+^ ion, which influence ligand binding and channel activity.

Interestingly, a structurally similar Cys_3_-His_1_ Zn^2+^-binding motif has recently been discovered in human TRPC5 (Wright et al., 2020). While being non-related in sequence to our motif identified in TRPM channels, this motif is also conserved across all TRPC channels, indicating that both TRPM as well as TRPC channels contain a conserved intracellular Zn^2+^-binding site.

## Materials and Methods

### Key Resources Table

| Reagent type (species)<br>or resource | Designation | Source or<br>reference | Identifiers |
| --- | --- | --- | --- |
| Gene | TRPM2 | GenScript | ODa47363D |
| Strain, strain background<br>(*E. coli*) | BL21 Gold (DE3) | Agilent | 230132 |
| Strain, strain background<br>(*E. coli*) | B834 (DE3) | Merck | 69041 |
| Chemical compound | L selenomethionine | Serva | 77765.01 |
| Chemical compound | ADPR | Sigma-Aldrich | A0752 |
| Chemical compound | 2'-deoxy-ADPR | BioLog | D 227-01 |
| Software, algorithm | XDS | (Kabsch, 2010) | http://xds.mpimf-heidelberg.mpg.de/ |
| Software, algorithm | AIMLESS | (Evans & Murshudov, 2013) | http://www.ccp4.ac.uk/html/aimless.html |
| Software, algorithm | SHELXCD | (Sheldrick, 2008) | https://shelx.uni-goettingen.de/ |
| Software, algorithm | ARP/wARP | (Langer et al., 2008) | https://www.embl-hamburg.de/ARP/ |
| Software, algorithm | Coot 0.8.9.1 | (Emsley et al., 2010) | https://www2.mrc-lmb.cam.ac.uk/personal/pemsley/coot/ |
| Software, algorithm | PHENIX 1.16 | (Liebschner et al., 2019) | https://www.phenix-online.org/ |
| Software, algorithm | Clustal Omega | (Madeira et al., 2019) | https://www.ebi.ac.uk/Tools/msa/clustalo/ |

### Materials

All chemicals were of analytical grade and obtained from Roth (Karlsruhe, Germany) or Sigma Aldrich / Merck (Darmstadt, Germany).

### Protein expression and purification

The sequence coding for the zebrafish TRPM2 MHR1/2 domain (drMHR1/2, residues 1-419 of drTRPM2) was cloned into a pnEK-vH vector bearing a TEV-cleavable N-terminal His_6_-tag. Protein production was carried out in *E. coli* BL21 Gold (DE3) cells in Terrific Broth (TB) media supplemented with 25 µg/mL kanamycin. Cells were grown at 37 °C until a cell density of 1 (measured at 600 nm) was reached. After induction with 0.1 mM isopropyl β-D-1-thiogalactopyranoside (IPTG), the cells were grown for further 16 hours at 20 °C. The production of selenomethionine-labelled protein was carried out in the methionine auxotroph *E. coli* strain B834 (Wood, 1966) in M9 minimal medium supplemented with 70 mg/L L-selenomethionine (Serva) and 25 µg/mL kanamycin. Cells were grown at 37 °C to an OD_600_ of 0.6 and after induction with 0.1 mM IPTG, the target protein was expressed for further 16 hours at 20 °C.

The cells were harvested by centrifugation (5,000×g for 25 min) and lysed in 25 mM Tris pH 7.5, 300 mM NaCl, 5% (v/v) glycerol, 5 mM 2-mercaptoethanol using a high-pressure homogenizer (EmulsiFlex-C3, Avestin). Cell debris was removed by centrifugation (39,000×g for 45 min) and the His_6_-tagged protein was purified from the supernatant using immobilized metal affinity chromatography (IMAC) with Ni-NTA resin (Roth). After His-tag removal by TEV protease (1/10 w:w) and reverse Ni-NTA purification, the target protein was further purified by gel filtration on a Superdex S200 increase 10/300 column using buffer M (for wild type protein, 25 mM HEPES pH 7.5, 150 mM NaCl, 5 mM CaCl_2_, 5 mM MgCl_2_) or buffer MS (for Se-Met-labelled protein, 25 mM HEPES pH 7.5, 150 mM NaCl, 5 mM CaCl_2_, 5 mM MgCl_2_, 5 mM 2-mercaptoethanol). Peak fractions were pooled and protein identity confirmed by SDS-PAGE and mass spectrometry.

### Isothermal titration calorimetry (ITC)

ITC measurements were carried out at 25°C using a MicroCal ITC-200 isothermal titration calorimeter (Malvern Panalytcal) and thermodynamic parameters were analyzed using the MicroCal ORIGIN™ software. The ligands ADPR and 2’-deoxy ADPR were each dissolved in buffer M (see purification) to a concentration of 500 µM and placed in the syringe. After an initial injection of 0.5 µL, 18 regular injections of 2 µL were added to 10 µM drMHR1/2 in the sample cell. The individual injections were interspaced by 150 s and stirring speed was set to 750 rpm. Heat of dilution was obtained by titrating ADPR into buffer M and baseline corrections were carried out accordingly. All ITC experiments were performed as triplicates and errors are reported as standard deviations of the mean K_D_ value.

### Differential scanning fluorimetry (nDSF)

nDSF measurements were carried out on a Prometheus NT.48 system (Nanotemper). 10 µM drMHR1/2 protein was mixed with varying amounts (600 nM -2 mM) of ADPR or 2’-deoxy ADPR in buffer M (see purification). Thermal unfolding was measured by following intrinsic tryptophan fluorescence during a thermal ramp (1°C/min). The PR.ThermControl software (Nanotemper) was used to determine melting temperatures. Binding parameter analysis was performed by simple Hill fit in the ORIGIN™ software.

### Crystallization

Crystals of selenomethionine-labelled drMHR1/2 protein were grown by sitting drop vapour diffusion technique. 1 µL of purified protein (4 mg/mL) was mixed with 1 µL of the precipitant mix (Tris, BICINE, Diethylene glycol, Triethylene glycol, Tetraethylene glycol, Pentaethylene glycol, glycerol, PEG4000, Jeffamine M-600). Crystals of triangular shape appeared after one to three days, reaching sizes of approximately 50-120 µm.

### Structure determination

X-ray diffraction data were collected at 100 K at the PETRA III/EMBL P14 beamline. All datasets were processed with XDS (Kabsch, 2010) and merged with AIMLESS (Evans & Murshudov, 2013). Heavy atom site identification and phasing was performed with SHELXCD (Sheldrick, 2008). A combination of ARP/wARP (Langer et al., 2008) and COOT (Emsley et al., 2010) was used for automatic and manual model building, respectively. Refinement was carried out in PHENIX (Liebschner et al., 2019). The final model corresponds to residues 38-423. All data collection and refinement statistics are summarized in Suppl. Table 1 (supplementary information).

### Multiple sequence alignment

Multiple sequence alignment of TRPM2 and homologues was performed using the Clustal Omega server (Madeira et al., 2019). Results were visualized with Jalview (https://www.jalview.org/).

### Cell surface biotinylation assay

Human embryonic kidney (HEK293) cells were grown in Dulbecco’s Modified Eagle Medium (DMEM) with 4.5 g/L Glucose and GlutaMAX™ (Gibco) at 37°C and 5% CO_2_. Full-length human TRPM2 variants (wild type, C89A, C91A, C89A C91A double mutant) were cloned into a pIRES2-EGFP vector allowing for fluorescent transfection control. Transfection was performed with Lipofectamine LTX (Thermo Fisher) according to the manufacturer’s instructions. Cells were grown for 48 h after transfection, washed with D-PBS (Gibco) and incubated with 1 mg/mL EZ-Link Sulfo-NHS-LC-Biotin (Thermo Fisher) in order to biotinylate cell surface proteins. After detachment with 2 mM EDTA in D-PBS, the cells were collected, centrifuged (500 g for 5 min) and washed with D-PBS. Cell lysis and membrane protein extraction was performed using the ProteoExtract Native Membrane kit (Merck Millipore) according to the manufacturer’s protocol. Solubilized membrane protein samples were quantified by Bradford assay.

NeutraVidin Agarose beads (70 µL, Thermo Fisher) were used to isolate the biotinylated cell surface proteins. The beads were incubated with total membrane protein samples (600 µg) for 18 h while rotating at 4°C. After washing with extraction buffer II from the kit mentioned above, the beads as well as the total membrane protein samples were mixed with SDS sample buffer (with 5% 2-mercapto ethanol) and heated to 75°C for 5 min.

The samples were analyzed by western blot. After SDS-PAGE (4-20% Protean precast gel, BioRad) and transfer to a PVDF membrane (Merck Millipore), the membrane was cut at 140 kDa in order to simultaneously detect TRPM2 (171 kDa) and the reference Na/K-ATPase (100 kDa). The membrane parts were probed for 18 h with anti-hsTRPM2 antibody from rabbit (Novus #nb500-241 at 1:50000 dilution) and anti-Na^+^/K^+^-ATPase antibody from rabbit (Cell Signaling #3010 at 1:1000 dilution), respectively. Both primary antibodies were detected with an HRP-conjugated anti-rabbit secondary antibody (Dianova #111-035-045 at 1:10000 dilution) for 1 h. Chemiluminescent detection was performed using the SuperSignal West Pico substrate (Thermo Fisher).

### Electrophysiology

Human embryonic kidney (HEK293) cells were grown in the conditions mentioned above and transfected with pIRES2-EGFP expression vectors containing hsTRPM2 variants (wild type, C89A, C91A, C89A C91A double mutant) 24 h before the patch clamp experiments. Transfection was performed with jetPEI reagent (PolyPlus Transfection). The transfection complex (5 µg DNA and 10 µL jetPEI) was incubated for 30 min before 2.5 × 10^5^ cells were added. The cell suspension was subsequently seeded to 35 mm dishes at low density to allow for single cell measurements.

The cell culture medium was replaced by bath solution (in mM: 140 *N*-methyl-D-glucamine (NMDG), 5 KCl, 3.3 MgCl_2_, 1 CaCl_2_,5 D-glucose, 10 HEPES, adjusted to pH 7.4 with HCl) directly before the experiment. Patch pipettes were pulled from thin-walled borosilicate glass capillaries (1.1 mm × 1.5 mm × 80 mm) with a Sutter P-97 horizontal puller and had a resistance between 1.5 MΩ and 2.5 MΩ. Pipettes were filled with the intracellular solution (in mM: 0.1 ADPR, 120 KCl, 8 NaCl, 1 MgCl_2_, 10 EGTA, 5.6 CaCl_2_, 10 HEPES, adjusted to pH 7.2 with KOH). Currents were recorded in whole cell mode using a HEKA EPC-10 amplifier and the PATCHMASTER software (HEKA Elektronik). TRPM2 activity was recorded using repetitive voltage ramps (−85 mV to +20 mV over 140 ms every 5 s for 450s). Cells were clamped to a holding potential of -50 mV before and in between ramps.

Statistical analysis was carried out with GRAPHPAD PRISM (v9.2.0; GraphPad Software) using one-way ANOVA. Dunnet’s multiple comparisons test was applied to compare the hsTRPM2 variants with the untransfected control.

## Acknowledgements

We are grateful to the staff at beamlines P14 and P13 (EMBL, Hamburg) and thank members of the Tidow and Fliegert labs for helpful discussions. We acknowledge access to the Sample Preparation and Characterization (SPC) Facility of EMBL, Hamburg. This work was supported by the Deutsche Forschungsgemeinschaft (DFG) (SFB1328, project A05 to HT and RF).

## Notes

The authors declare no competing financial interest.

## Footnotes

The abbreviations used are: ADPR, adenosine diphosphate ribose; DMEM, Dulbecco’s Modified Eagle Medium; EDTA, ethylenediaminetetraacetic acid; EM, electron microscopy; IMAC, immobilized metal affinity chromatography; IPTG, isopropyl β-D-1-thiogalactopyranoside; ITC, isothermal titration calorimetry; MHR, TRPM homology region; NAD, nicotinamide adenine dinucleotide; Ni-NTA, Ni-nitrilotriacetic acid; NMDG, *N*-methyl-D-glucamine; PARG, poly-ADPR glycohydrolase; PARP, poly-ADPR polymerase; ROS, reactive oxygen species; SAD, single-wavelength anomalous diffraction; SDS-PAGE, sodium dodecyl sulphate-polyacrylamide gel electrophoresis; SD, standard deviation; TEV, tobacco etch virus; TRPM, transient receptor potential melastatin.

## Data Availability

Structural coordinates and structural factors have been deposited in the RCSB Protein Data Bank under accession number 7AOV (see Suppl. Table 1).

## Supplementary Information

### Supplementary Tables

**Suppl. Table 1.** Data collection and refinement statistics. Data from the last resolution shell are in parentheses.

|  | **drMHR1/2: PDB ID 7AOV** |
| --- | --- |
| **Data collection** |  |
| Beamline | PETRA III, P13 |
| Wavelength (Å) | 0.98 |
| Resolution (Å) | 46.6 - 2.0 (2.07 - 2.00) |
| Space group | P 1 2<sub>1</sub> 1 |
| Cell dimensions |  |
| a, b, c (Å) | 47.8, 188.1, 47.8 |
| α, β, γ (°) | 90.0, 102.7, 90.0 |
| Completeness (%) | 97.2 (96.9) |
| Mean I/σ (I) | 10.3 (1.5) |
| R<sub>meas</sub> | 0.288 (1.88) |
| R<sub>pim</sub> | 0.077 (0.51) |
| CC<sub>1/2</sub> | 0.996 (0.552) |
| Multiplicity | 13.9 (13.6) |
| **Refinement** |  |
| Reflections used in refinement | 53670 (5367) |
| R<sub>work</sub> / R<sub>free</sub> | 0.19 / 0.23 |
| Number of non-hydrogen atoms | 6373 |
| macromolecules | 5956 |
| ligands/ions | 2 |
| Average B-factor (Å<sup>2</sup>) | 45.7 |
| macromolecules (Å<sup>2</sup>) | 45.5 |
| ligands/ions (Å<sup>2</sup>) | 30.5 |
| RMS<sub>bonds</sub> (Å) | 0.008 |
| RMS<sub>angles</sub> (°) | 0.97 |
| Ramachandran |  |
| Favored regions (%) | 95.8 |
| Allowed regions (%) | 3.8 |
| Outliers (%) | 0.4 |
| Rotamer outliers (%) | 0.0 |

### Supplementary Figures

**Suppl. Fig. S1.**
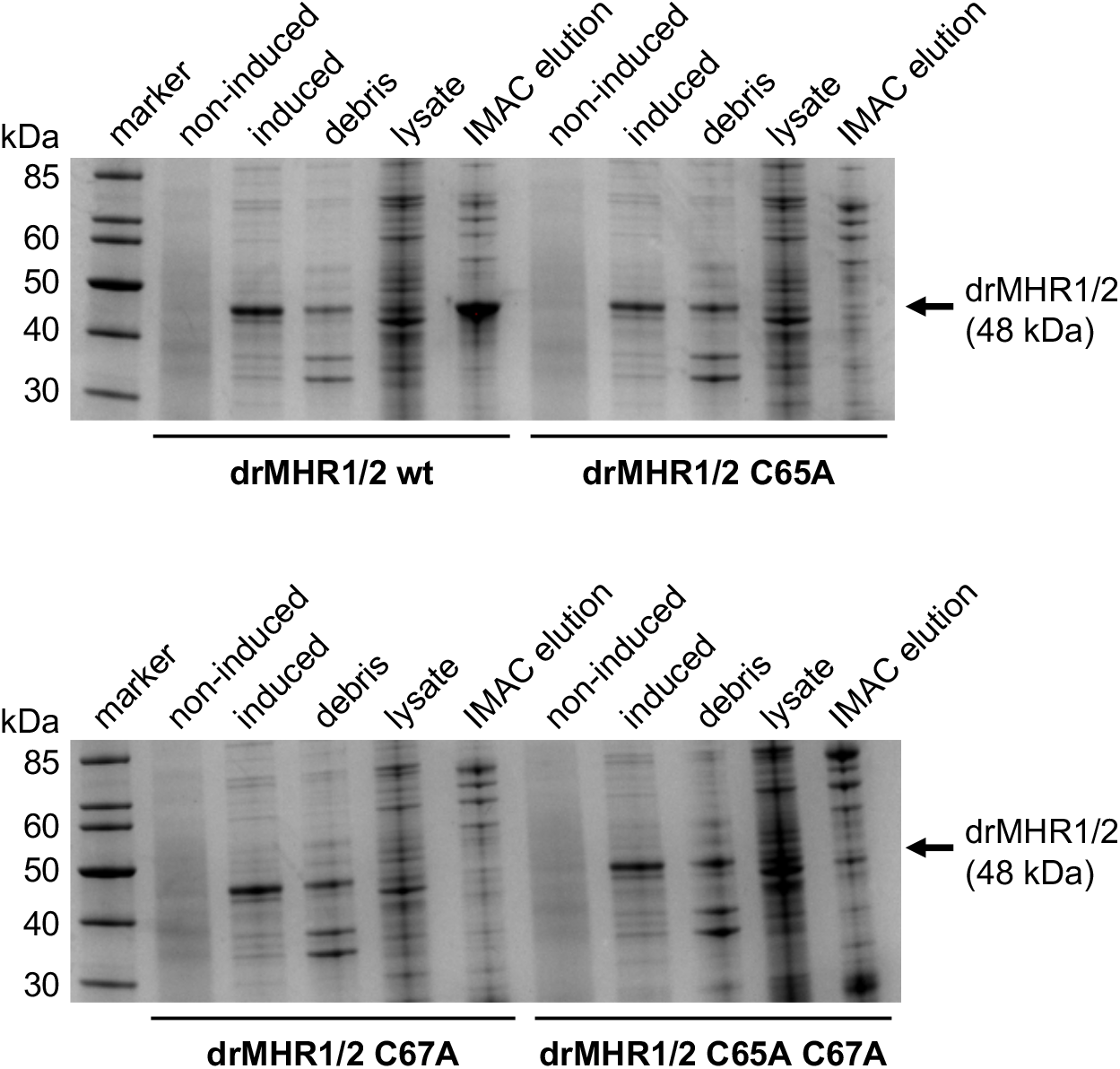
Mutations within the Zn^2+^-binding domain prove its importance for protein integrity. Coomassie-stained SDS-PAGE gels show that drMHR1/2 mutants (C65A, C67A and C65A C67A double mutant) are expressed like the wild-type (induced cells lane) but do not seem to be soluble (missing drMHR1/2 band in lysate and IMAC elution lanes).

## References

Abriel, H., Syam, N., Sottas, V., Amarouch, M. Y., & Rougier, J.-S. (2012). TRPM4 channels in the cardiovascular system: Physiology, pathophysiology, and pharmacology. Biochemical Pharmacology, 84(7), 873–881. https://doi.org/10.1016/j.bcp.2012.06.021

Abuarab, N., Munsey, T. S., Jiang, L. H., Li, J., & Sivaprasadarao, A. (2017). High glucose-induced ROS activates TRPM2 to trigger lysosomal membrane permeabilization and Zn2+-mediated mitochondrial fission. Science Signaling, 10(490), 1–12. https://doi.org/10.1126/scisignal.aal4161

Bouron, A., & Oberwinkler, J. (2014). Contribution of calcium-conducting channels to the transport of zinc ions. Pflugers Archiv European Journal of Physiology, 466(3), 381– 387. https://doi.org/10.1007/s00424-013-1295-z

Buelow, B., Song, Y., & Scharenberg, A. M. (2008). The Poly(ADP-ribose) polymerase PARP-1 is required for oxidative stress-induced TRPM2 activation in lymphocytes. The Journal of Biological Chemistry, 283(36), 24571–24583. https://doi.org/10.1074/jbc.M802673200

Emsley, P., Lohkamp, B., Scott, W. G., & Cowtan, K. (2010). Features and development of Coot. Acta Crystallographica Section D: Biological Crystallography, 66(4), 486–501. https://doi.org/10.1107/S0907444910007493

Evans, P. R., & Murshudov, G. N. (2013). How good are my data and what is the resolution? Acta Crystallographica Section D: Biological Crystallography, 69(7), 1204–1214. https://doi.org/10.1107/S0907444913000061

Fliegert, R., Bauche, A., Wolf Pérez, A. M., Watt, J. M., Rozewitz, M. D., Winzer, R., Janus, M., Gu, F., Rosche, A., Harneit, A., Flato, M., Moreau, C., Kirchberger, T., Wolters, V., Potter, B. V. L., & Guse, A. H. (2017). 2′-Deoxyadenosine 5′-diphosphoribose is an endogenous TRPM2 superagonist. Nature Chemical Biology, 13(9), 1036–1044. https://doi.org/10.1038/nchembio.2415

Fukada, T., Yamasaki, S., Nishida, K., Murakami, M., & Hirano, T. (2011). Zinc homeostasis and signaling in health and diseases. Journal of Biological Inorganic Chemistry, 16(7), 1123–1134. https://doi.org/10.1007/s00775-011-0797-4

Granzotto, A., Canzoniero, L. M. T., & Sensi, S. L. (2020). A Neurotoxic Ménage-à-trois: Glutamate, Calcium, and Zinc in the Excitotoxic Cascade. Frontiers in Molecular Neuroscience, 13(November), 1–13. https://doi.org/10.3389/fnmol.2020.600089

Hao, Q., & Maret, W. (2006). Aldehydes release zinc from proteins. A pathway from oxidative stress/lipid peroxidation to cellular functions of zinc. FEBS Journal, 273(18), 4300–4310. https://doi.org/10.1111/j.1742-4658.2006.05428.x

Hecquet, C. M., Zhang, M., Mittal, M., Vogel, S. M., Di, A., Gao, X., Bonini, M. G., & Malik, A. B. (2014). Cooperative interaction of trp melastatin channel transient receptor potential (TRPM2) with its splice variant TRPM2 short variant is essential for Endothelial cell Apoptosis. Circulation Research, 114(3), 469–479. https://doi.org/10.1161/CIRCRESAHA.114.302414

Howard, M., Grimaldi, J., Bazan, J., Lund, F., Santos-Argumedo, L., Parkhouse, R., Walseth, T., & Lee, H. (1993). Formation and hydrolysis of cyclic ADP-ribose catalyzed by lymphocyte antigen CD38. Science, 262(5136), 1056–1059. https://doi.org/10.1126/science.8235624

Huang, Y., Fliegert, R., Guse, A. H., Lü, W., & Du, J. (2020). A structural overview of the ion channels of the TRPM family. Cell Calcium, 85(September 2019), 102111. https://doi.org/10.1016/j.ceca.2019.102111

Huang, Y., Roth, B., Lü, W., & Du, J. (2019). Ligand recognition and gating mechanism through three ligand-binding sites of human TRPM2 channel. ELife, 8, 1–18. https://doi.org/10.7554/eLife.50175

Huang, Y., Winkler, P. A., Sun, W., Lü, W., & Du, J. (2018). Architecture of the TRPM2 channel and its activation mechanism by ADP-ribose and calcium. Nature, 562(7725), 145–149. https://doi.org/10.1038/s41586-018-0558-4

Iordanov, I., Mihályi, C., Tóth, B., & Csanády, L. (2016). The proposed channel-enzyme transient receptor potential melastatin 2 does not possess ADP ribose hydrolase activity. ELife, 5(JULY), 1–20. https://doi.org/10.7554/eLife.17600

Jimenez, I., Prado, Y., Marchant, F., Otero, C., Eltit, F., Cabello-Verrugio, C., Cerda, O., & Simon, F. (2020). TRPM Channels in Human Diseases. Cells, 9(12). https://doi.org/10.3390/cells9122604

Kabsch, W. (2010). Xds. Acta Crystallographica. Section D, Biological Crystallography, 66(Pt 2), 125–132. https://doi.org/10.1107/S0907444909047337

Kambe, T., Tsuji, T., Hashimoto, A., & Itsumura, N. (2015). The physiological, biochemical, and molecular roles of zinc transporters in zinc homeostasis and metabolism. Physiological Reviews, 95(3), 749–784. https://doi.org/10.1152/physrev.00035.2014

Knowles, H., Heizer, J. W., Li, Y., Chapman, K., Ogden, C. A., Andreasen, K., Shapland, E., Kucera, G., Mogan, J., Humann, J., Lenz, L. L., Morrison, A. D., & Perraud, A.-L. (2011). Transient Receptor Potential Melastatin 2 (TRPM2) ion channel is required for innate immunity against Listeria monocytogenes. Proceedings of the National Academy of Sciences, 108(28), 11578–11583. https://doi.org/10.1073/pnas.1010678108

Langer, G., Cohen, S. X., Lamzin, V. S., & Perrakis, A. (2008). Automated macromolecular model building for X-ray crystallography using ARP/wARP version 7. Nature Protocols, 3(7), 1171–1179. https://doi.org/10.1038/nprot.2008.91

Liebschner, D., Afonine, P. V., Baker, M. L., Bunkoczi, G., Chen, V. B., Croll, T. I., Hintze, B., Hung, L. W., Jain, S., McCoy, A. J., Moriarty, N. W., Oeffner, R. D., Poon, B. K., Prisant, M. G., Read, R. J., Richardson, J. S., Richardson, D. C., Sammito, M. D., Sobolev, O. V., … Adams, P. D. (2019). Macromolecular structure determination using X-rays, neutrons and electrons: Recent developments in Phenix. Acta Crystallographica Section D: Structural Biology, 75, 861–877. https://doi.org/10.1107/S2059798319011471

Madeira, F., Park, Y. M., Lee, J., Buso, N., Gur, T., Madhusoodanan, N., Basutkar, P., Tivey, A. R. N., Potter, S. C., Finn, R. D., & Lopez, R. (2019). The EMBL-EBI search and sequence analysis tools APIs in 2019. Nucleic Acids Research, 47(W1), W636–W641. https://doi.org/10.1093/nar/gkz268

McKemy, D. D., Neuhausser, W. M., & Julius, D. (2002). Identification of a cold receptor reveals a general role for TRP channels in thermosensation. Nature, 416(6876), 52–58. https://doi.org/10.1038/nature719

McNulty, S., & Fonfria, E. (2005). The role of TRPM channels in cell death. Pflugers Archiv European Journal of Physiology, 451(1), 235–242. https://doi.org/10.1007/s00424-005-1440-4

Montell, C. (2003). Mg2+ Homeostasis: The Mg2+nificent TRPM Chanzymes. Current Biology, 13(20), 799–801. https://doi.org/10.1016/j.cub.2003.09.048

Mortadza, S. S., Sim, J. A., Stacey, M., & Jiang, L. H. (2017). Signalling mechanisms mediating Zn 2+ -induced TRPM2 channel activation and cell death in microglial cells. Scientific Reports, 7(February), 1–15. https://doi.org/10.1038/srep45032

Nikiforov, A., Kulikova, V., & Ziegler, M. (2015). The human NAD metabolome: Functions, metabolism and compartmentalization. Critical Reviews in Biochemistry and Molecular Biology, 50(4), 284–297. https://doi.org/10.3109/10409238.2015.1028612

Park, S. E., Song, J. H., Hong, C., Kim, D. E., Sul, J.-W., Kim, T.-Y., Seo, B.-R., So, I., Kim, S.-Y., Bae, D.-J., Park, M.-H., Lim, H. M., Baek, I.-J., Riccio, A., Lee, J.-Y., Shim, W. H., Park, B., Koh, J.-Y., & Hwang, J. J. (2019). Contribution of Zinc-Dependent Delayed Calcium Influx via TRPC5 in Oxidative Neuronal Death and its Prevention by Novel TRPC Antagonist. Molecular Neurobiology, 56(4), 2836–2837. https://doi.org/10.1007/s12035-018-1447-4

Pérez, C. A., Huang, L., Rong, M., Kozak, J. A., Preuss, A. K., Zhang, H., Max, M., & Margolskee, R. F. (2002). A transient receptor potential channel expressed in taste receptor cells. Nature Neuroscience, 5(11), 1169–1176. https://doi.org/10.1038/nn952

Perraud, A. L., Fleig, A., Dunn, C. A., Bagley, L. A., Launay, P., Schmitz, C., Stokes, A. J., Zhu, Q., Bessman, M. J., Penner, R., Kinet, J. P., & Scharenberg, A. M. (2001). ADP-ribose gating of the calcium-permeable LTRPC2 channel revealed by Nudix motif homology. Nature, 411(6837), 595–599. https://doi.org/10.1038/35079100

Schmitz, C., & Perraud, A.-L. (2005). The TRPM cation channels in the immune context. Current Pharmaceutical Design, 11(21), 2765–2778. https://doi.org/10.2174/1381612054546851

Sheldrick, G. M. (2008). A short history of SHELX. Acta Crystallographica Section A: Foundations of Crystallography, 64(1), 112–122. https://doi.org/10.1107/S0108767307043930

Siemens, J., Pohle, J., Wang, H., Song, K., Wende, H., Heppenstall, P., Reis, F. d. C., & Kamm, G. B. (2016). The TRPM2 channel is a hypothalamic heat sensor that limits fever and can drive hypothermia. Science, 353(6306), 1393–1398. https://doi.org/10.1126/science.aaf7537

Simon, F., Varela, D., & Cabello-Verrugio, C. (2013). Oxidative stress-modulated TRPM ion channels in cell dysfunction and pathological conditions in humans. Cellular Signalling, 25(7), 1614–1624. https://doi.org/10.1016/j.cellsig.2013.03.023

Sun, Y., Sukumaran, P., Schaar, A., & Singh, B. B. (2015). TRPM7 and its role in neurodegenerative diseases. Channels, 9(5), 253–261. https://doi.org/10.1080/19336950.2015.1075675

Tan, C. H., & McNaughton, P. A. (2016). The TRPM2 ion channel is required for sensitivity to warmth. Nature, 536(7617), 460–463. https://doi.org/10.1038/nature19074

Tóth, B., Iordanov, I., & Csanády, L. (2020). Selective profiling of N-and C-terminal nucleotide-binding sites in a TRPM2 channel. The Journal of General Physiology, 152(5), 1–13. https://doi.org/10.1085/jgp.201912533

Vennekens, R., Mesuere, M., & Philippaert, K. (2018). TRPM5 in the battle against diabetes and obesity. Acta Physiologica, 222(2), e12949. https://doi.org/10.1111/apha.12949

Wang, L., Fu, T.-M., Zhou, Y., Xia, S., Greka, A., & Wu, H. (2018). Structures and gating mechanism of human TRPM2. Science (New York, N.Y.), 362(6421), eaav4809. https://doi.org/10.1126/science.aav4809

Wood, W. B. (1966). Host specificity of DNA produced by Escherichia coli: Bacterial mutations affecting the restriction and modification of DNA. Journal of Molecular Biology, 16(1), 118–133. https://doi.org/10.1016/S0022-2836(66)80267-X

Wright, D. J., Simmons, K. J., Johnson, R. M., Beech, D. J., Muench, S. P., & Bon, R. S. (2020). Human TRPC5 structures reveal interaction of a xanthine-based TRPC1/4/5 inhibitor with a conserved lipid binding site. Communications Biology, 3(1), 1–11. https://doi.org/10.1038/s42003-020-01437-8

Yang, W., Manna, P. T., Zou, J., Luo, J., Beech, D. J., Sivaprasadarao, A., & Jiang, L. H. (2011). Zinc inactivates melastatin transient receptor potential 2 channels via the outer pore. Journal of Biological Chemistry, 286(27), 23789–23798. https://doi.org/10.1074/jbc.M111.247478

Ye, M., Yang, W., Ainscough, J. F., Hu, X.-P., Li, X., Sedo, A., Zhang, X.-H., Zhang, X., Chen, Z., Li, X.-M., Beech, D. J., Sivaprasadarao, A., Luo, J.-H., & Jiang, L.-H. (2014). TRPM2 channel deficiency prevents delayed cytosolic Zn2+ accumulation and CA1 pyramidal neuronal death after transient global ischemia. Cell Death & Disease, 5(11), e1541. https://doi.org/10.1038/cddis.2014.494

Yin, Y., Wu, M., Hsu, A. L., Borschel, W. F., Borgnia, M. J., Lander, G. C., & Lee, S.-Y. (2019). Visualizing structural transitions of ligand-dependent gating of the TRPM2 channel. Nature Communications, 10(1), 3740. https://doi.org/10.1038/s41467-019-11733-5

Yu, P., Wang, Q., Zhang, L. H., Lee, H. C., Zhang, L., & Yue, J. (2012). A Cell Permeable NPE Caged ADP-Ribose for Studying TRPM2. PLoS ONE, 7(12). https://doi.org/10.1371/journal.pone.0051028

Zheng, H., Cooper, D. R., Porebski, P. J., Shabalin, I. G., Handing, K. B., & Minor, W. (2017). CheckMyMetal: a macromolecular metal-binding validation tool. Acta Crystallographica. Section D, Structural Biology, 73(Pt 3), 223–233. https://doi.org/10.1107/S2059798317001061

Zocchi, E., Franco, L., Guida, L., Benatti, U., Bargellesi, A., Malavasi, F., Lee, H. C., & Deflora, A. (1993). A Single Protein Immunologically Identified as CD38 Displays NAD+ Glycohydrolase, ADP-Ribosyl Cyclase and Cyclic ADP-Ribose Hydrolase Activities at the Outer Surface of Human Erythrocytes. Biochemical and Biophysical Research Communications, 196(3), 1459–1465. https://doi.org/10.1006/bbrc.1993.2416

